# Post-transcriptional regulation of the MiaA prenyl transferase by CsrA and the small RNA CsrB in *E. coli*

**DOI:** 10.1101/2024.02.28.582573

**Authors:** Joseph I Aubee, Kinlyn Williams, Alexandria Adigun, Olufolakemi Olusanya, Jalisa Nurse, Karl M Thompson

## Abstract

To determine the role of small RNAs (sRNAs) in the regulation of *miaA*, we constructed a chromosomal *miaA*-*lacZ* translational fusion driven by the arabinose responsive P_BAD_ promoter and used it to screen against an *Escherichia coli* small RNA library (containing small RNAs driven by the IPTG inducible P*_Lac_* promoter). Our genetic screen and quantitative β-galactosidase assays identified CsrB and its cognate protein CsrA as potential regulators of *miaA* expression in *Escherichia coli*. Consistent with our hypothesis that CsrA regulates MiaA post-transcriptional gene expression through binding to the MiaA mRNA 5’ UTR, and CsrB binds and regulates MiaA post-transcriptional gene expression through sequestration of CsrA levels, a deletion of *csrA* significantly reduced expression of the reporter fusion as well as reducing *miaA* mRNA levels. These results suggest under conditions where CsrA is inhibited, MiaA translation and thus MiaA-dependent tRNA modification may be limiting.

**IMPORTANCE:** We previously demonstrated a role for the i^6^A modification in the tuning of transcripts for several stress response genes in *E. coli*. The i^6^A tRNA modification is catalyzed by the tRNA prenyl transferase encoded by the *miaA* gene. We set out to identify posttranscriptional regulators of the enzyme necessary for the catalysis of i^6^A, MiaA, to further understand factors influencing i^6^A levels in the cell. We identified the CsrA RNA Binding Protein, the CsrB Small RNA, and RNA Degradosome enzymes: RNaseE and PNPase as regulators of *miaA* expressioin at the post-transcriptional level. Identifying these post-transcripitonal regulators of *miaA* will help us understand factors influencing i^6^A levels and may guide future investigations into RNA modifications with regulatory effects on the transcriptome.

## INTRODUCTION

MiaA is a tRNA Isopentyl Transferase that catalyzes the prenylation of adenine 37 in the anticodon stem loop of tRNAs that read codons beginning with uridine (1–3). The resulting N^6^-(isopentenyl) adenosine 37 (i^6^A37) is the precursor for subsequent methylthiolation by the MiaB enzyme, resulting in the 2-methylthio-N^6^-(isopentenyl) adenosine (ms^2^i^6^A37) in *E. coli* (1–3). The function of MiaA has been the subject of study for several decades (1–7). Previous studies demonstrated a role for i^6^A37, and other RNA modifications, in translational fidelity by preventing translational aberrations such as ribosome pausing and ribosomal frameshifting (8–15). In addition, *E. coli miaA* mutants decrease cellular growth rates and promote spontaneous mutants, specifically GC – TA transversions (2, 3, 16). MiaA is highly conserved with homologues in both prokaryotes and eukaryotes (17–19). MiaA levels influence translational frameshifting to alter the global proteome, fitness, and virulence potential of Extraintestinal Pathogenic *E. coli* (ExPEC) (20). MiaA promotes virulence in *Shigella flexneri* (21). *Acinetobacter baumannii miaA* mutations exhibit Colistin resistance (22). *Streptomyces albus* requires *miaA* for proper morphological development and metabolic regulation (23, 24).

We previously identified MiaA as a regulatory factor necessary for the expression of the stationary phase and general stress response sigma factor RpoS in *E. coli* K12 (σ^S^) (25). MiaA affects the expression of two additional stress response genes in *E. coli* K12: Hfq and IraP, which are also involved in the regulation of RpoS (26, 27). The MiaA (i^6^A) sensitive genes identified in *E. coli* K12 thus far, have higher UUX-Leucine codon usage than the average genome wide UUX-Leucine codon usage (25, 27). MiaA promotes the expression of its targets by promoting efficient UUX-Leucine decoding (27). While these studies have expanded our understanding of i^6^A modification function, we still do not know the physiological or metabolic conditions that control i^6^A levels in *E. coli*. We reasoned that further characterization of MiaA synthesis would assist us in understanding the regulation of i^6^A levels in the cell and we sought to do that in this work.

The *miaA* gene is contained in a complex operon, immediately upstream of *hfq*, the gene encoding the RNA chaperone and host factor for Bacteriophage Qβ replication (28–30). The transcription of *miaA* is driven by two promoters, one of which is a heat shock promoter (*miaA*_P2(hs)_) that is recognized by the heat shock responsive alternative sigma factor, σ^32^ (30, 31). However, we know much less about the post-transcriptional regulatory factors that influence *miaA* expression. There are several clues in the literature that point to potential post-transcriptional regulation of MiaA. First, the transcript driven by the *miaA*_P2(h)_ promoter has a 270 nucleotide 5’ untranslated region (UTR) (30, 31). Long 5’ UTRs often associated with post-transcriptional regulatory processes. Second, a null mutation in *hfq*, resulted in elevated levels of multiple transcripts from the *miaA* superoperon (32). Third, MiaA transcript levels are increased in the absence of RNaseE and / or RNaseIII (31). RnaseE is an endoribonuclease that is essential for growth, and works with the 3’ to 5’ exoribonuclease Polynucleotide phosphorylase (PNPase), to process rRNA and tRNA and execiute the coordinated turnover of sRNAs and their mRNA targets (33–36). These previous results support the idea that MiaA expression is regulated at the post-transcriptional level. Post-transcripitonal regulation of *miaA* by Hfq and RNase would likely be mediated by small RNAs. Yet, prior to this work, no small RNA regulators of *miaA* have been identified. To close this gap, we executed a targeted screen of a pladmid sRNA library on a P*_BAD_*-*miaA27_P2_*-*lacZ* translational fusion strain, as previously described (37–39). We identified several candidate sRNA repressors of the *miaA*_P2(hs)_ transcript including SdsR (RyeB), ArcZ, GcvB, Spot42, and CsrB. While SdsR and CsrB had the most dramatic inhibitory effect of all candidate sRNAs regulators of the *miaA*_P2(hs)_ transcript in our genetic screen, the CsrB effect on *miaA* expression was the focus of subsequent experiments for this study.

CsrB is a small RNA that primarily acts to sequester the activity of the RNA binding protein CsrA (40). CsrA is a pleiotropic regulator of carbon metabolism and global RNA binding protein involved in direct post-transcriptional regulation of gene expression in *E. coli*, following binding to the 5’ untranslated regions (41–43). CsrA regulates the expression of *pgaABCD* and *flhDC* operons to influence biofilm formation and motility / flagellar synthesis, respectively, in *E. coli* (44, 45). CsrA binds to several mRNA transcripts and subsequently regulates the expression of these genes in *E. coli*. The regulatory processes, metabolic impact, and virulence promoting activities of CsrA have been identified in many other bacteria including *Escherichia coli*, *Salmonella Typhimurium*, *Pseudomonas sp*, *Serratia sp.*, *Campylobacter jejuni*, *Vibrio cholera*, *Yersinia pseudotuberculosis*, *Erwinia amylovora*, *Legionella pneumophila*, *Bacillus subtilis*, *Staphylococcus aureus*, *Clostridiodes difficile*, and *Acinetobacter baumannii* (46–61). CsrA activity was previously demonstrated to be regulated through sequestration by small RNAs CsrB and CsrC. However, recent studies have demonstrated an expanded number of 10 direct CsrA sRNA binding partners in vitro (62). Four of them bind to CsrA in vivo (62). In addition, recent studies suggest that CsrA binds sRNAs and promotes sRNA-mRNA complex formation, placing it in the category of RNA chaperones such as Hfq and ProQ (57, 63, 64). Since CsrA works with CsrB to regulate gene expression, we tested the effect of CsrA on MiaA expression. We show that CsrA is necessary for *miaA*_P2(hs)_ translation. We further extended the prior work on RNases and MiaA expression, demonstrating that RNaseE and PNPase from the Degradosome contribute to stabilization of the *miaA* mRNA transcript.

## MATERIALS AND METHODS

### Strains and Plasmids

All strains are derivatives of *E. coli* K12 MG1655 and are listed in Table 1. All plasmids used in this study are also listed in Table 1.

**Table 1.-.**
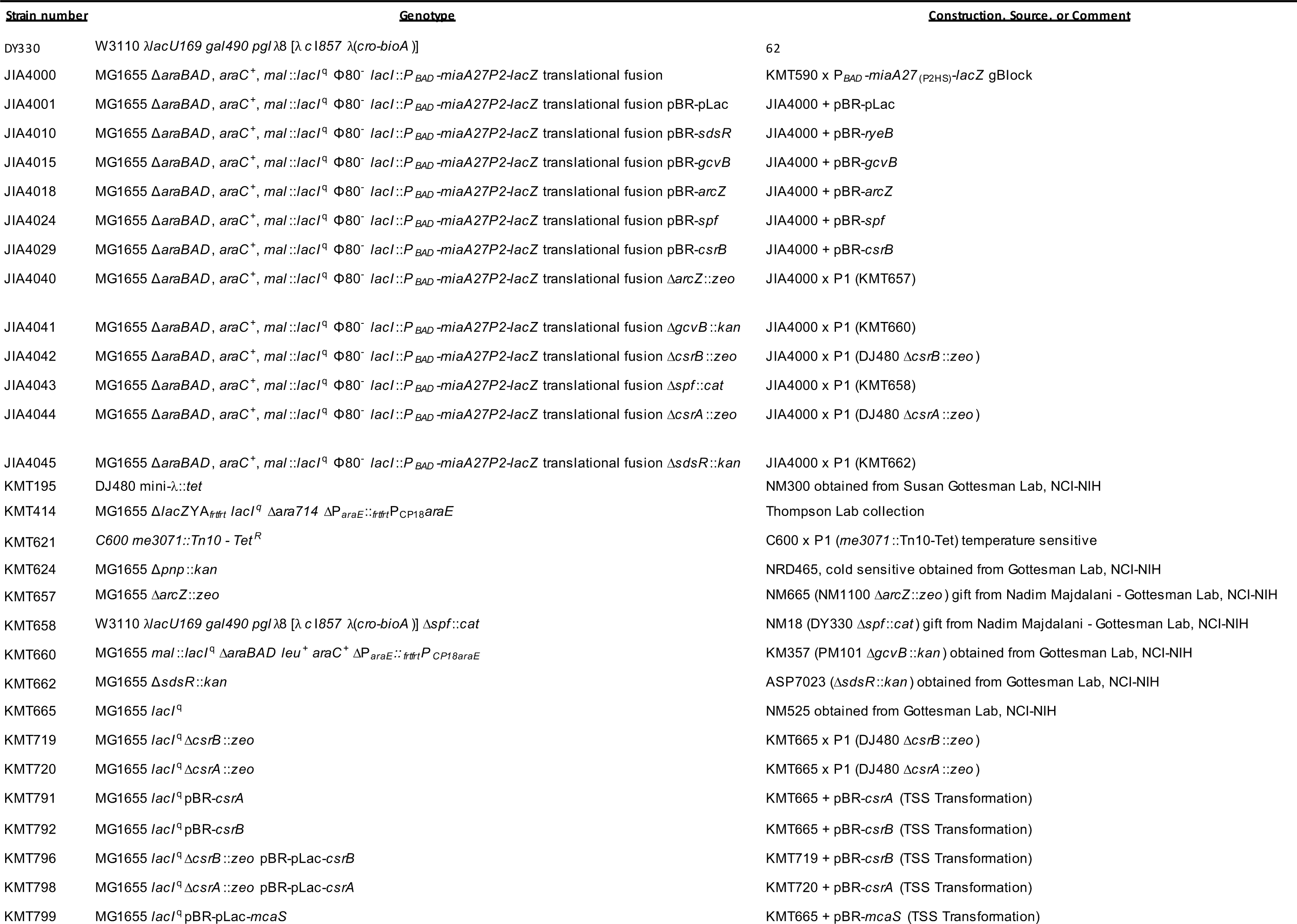
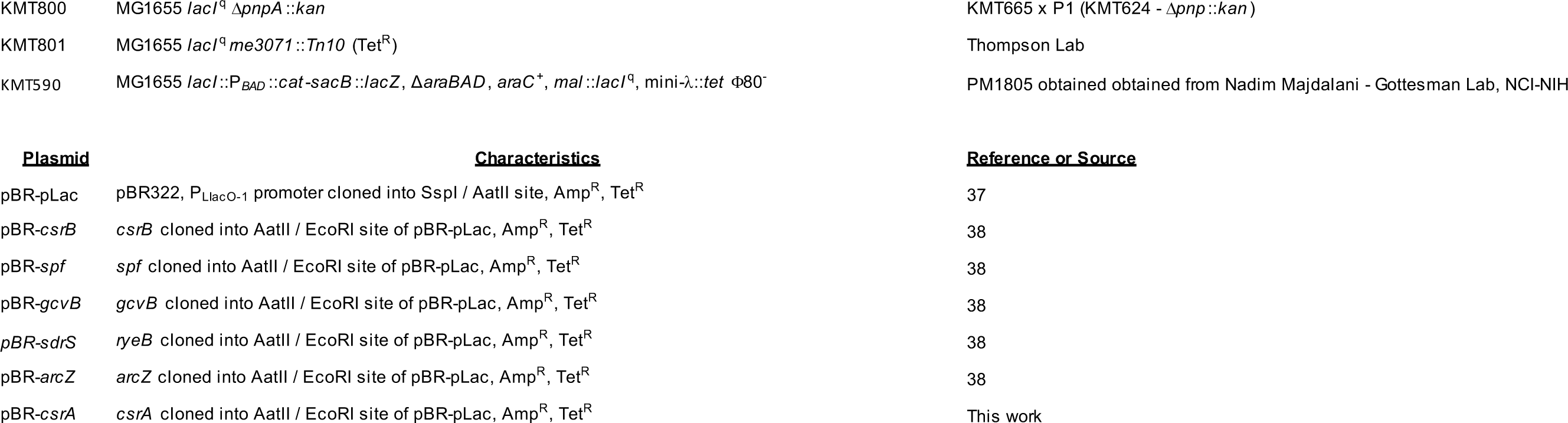
Strains and Plasmids.

### Media and Growth Conditions

*E. coli* strains were grown in Luria Bertani (LB) (Lennox) liquid media in a WS27 shaking water bath (ShelLab) for the experiments in this study. Recombinants for Lambda (λ)-Red based mutagenesis were selected for on LB agar plates supplemented with zeomycin to a final concentration of 25-50 μg/mL (LB-Zeo). Transductants, of *pnpA*::*kan* or *rne-3071 zce-726*::Tn*10* (*tet*) mutants, were grown on LB agar plates supplemented with either tetracycline or kanamycin to a final concentration of 25 μg/mL (LB-Tet or LB-Kan). The *rne-3071 zce-726*::Tn*10* mutant is temperature sensitive and requires growth at 30°C. To determine the effect of RNaseE on MiaA mRNA levels, *rne-3071 zce-726*::Tn*10* cultures were grown at 30°C and then shifted to 43.5°C to induce an RNaseE^−^ phenotype, as previously described (36). We then isolated total RNA for the analysis of MiaA mRNA levels or turnover using northern blot analysis. Strains carrying plasmids were grown in LB media supplemented with ampicillin to a final concentration of 100 μg/mL (LB-Amp) or on LB-Amp agar plates. To stimulate the expression of arabinose inducible fusion strains, cultures were first grown in LB or LB-Amp, supplemented with glucose to a final concentration of 0.2%, harvested by centrifugation, washed once with LB media, and resuspended in LB supplemented with arabinose to a final concentration of 0.2%. Small RNA plasmid library transformants were assayed on MacConkey-Lactose (Mac-Lac) agar plates supplemented with ampicillin to a final concentration of 100 µg/mL (Mac-Lac-Amp) to screen for changes in the Lactose phenotype of individual colonies following overnight growth at 37°C.

### General Molecular Biology Techniques

Plasmid DNA was isolated using The Column-Pure^TM^ Plasmid Mini-Prep Kit (Lamda Biotech), according to manufacturer instructions. Genomic DNA used for PCR reactions was isolated using The Column-Pure^TM^ Bacterial Genomic DNA Kit (Lamda Biotech), according to the manufacturer’s instructions. PCR reactions were performed to amplify allelic exchange substrates for mutagenesis. PCR amplification was done using the Taq Plus 2X PCR MasterMix (Lamda Biotech) with standard cycle conditions according to the manufacturer’s instructions. The annealing temperature for each PCR reaction was optimized to the T_m_ of the primers used (Table 2). All PCR reactions were purified using The Column-Pure^TM^ Clean-Up Kit (Lamda Biotech), according to the manufacturer’s instructions (Lamda Biotech).

**Table 2-.**
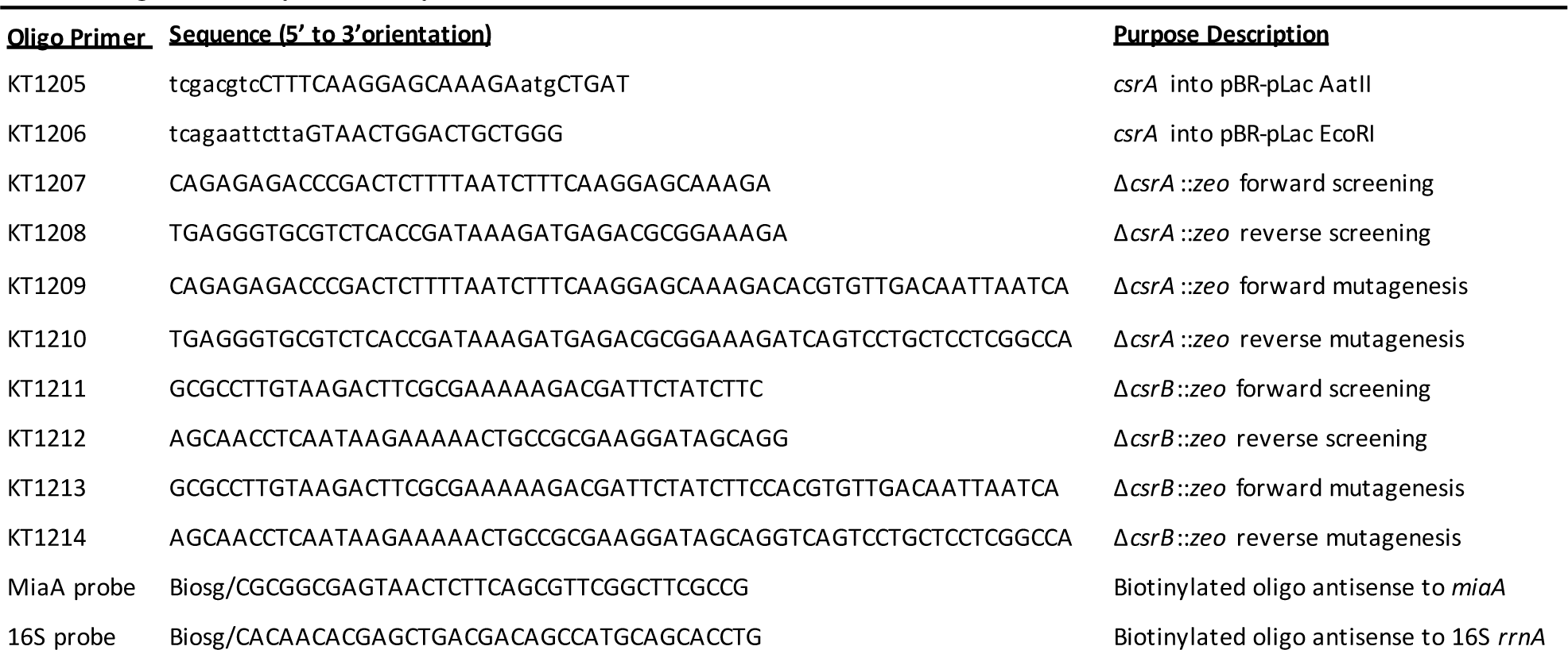
Oligonucleotide primers and probes.

### Genetic engineering and strain construction

Chromosomal mutagenesis was executed via Bacteriophage λ based recombineering and Bacteriophage P1 transduction as previously described and also outlined below (65, 66). Plasmids were moved using chemical transformation as described below.

#### Insertional Inactivation Mutagenesis using Recombineering

Deletion insertion mutations of *csrA*, *csrB*, and *miaA* were constructed using recombineering as previously described (67, 68). Briefly, a zeomycin resistance cassette with 50 bp of flanking homology to *csrA*, *csrB*, or *miaA* was synthesized by PCR to create an allelic exchange substrate for recombineering based mutagenesis. Then, the DJ480 mini-λ::*tet* strain was induced using heat shock and made electrocompetent with ice-cold water washes (67, 68). The allelic exchange substrates were then electroporated into the prepared cells, allowed to recover, and plated on LB-Zeo selectable media. Recombinants were confirmed to have the mutant using PCR and Sanger Sequencing.

#### P1 Transduction to move mutants between strains

The newly constructed Δ*csrA*::*zeo*, Δ*csrB*::*zeo*, Δ*miaA*::*zeo* mutations, as well as previously constructed *pnpA*::*kan* and *rne-3071 zce-726*::Tn*10* (*tet*) mutants were moved from the mini-λ::tet containing strain into clean genetic backgrounds (MG1655 or P*_BAD_*-*miaA27*_P2_-*lacZ* translational fusion strain) via generalized transduction using Bacteriophage P1 as previously described (69).

#### Chemical / Heat Shock Transformation to induce plasmid uptake

All Plasmids, including those from the small RNA library, were transformed into *E. coli* strains using Transformation Storage Solution (TSS) media and its associated transformation method as previously described (70). Briefly, the *E. coli* K-12 MG1655 recipient strain was grown in nutrient-rich Lennox Broth (LB) to OD_600_ 0.5. Upon reaching OD_600_ 0.5, the cells were harvested by centrifugation. The supernatant was then decanted, and the pellet re-suspended in 1/10th volume of ice-cold TSS media. 2μL (100ng) of plasmid DNA (pBR-pLac, pBR-pLac-*csrB*, pBR-pLac-*sdsR*, pBR-pLac-*spf*, pBR-pLac-*arcZ*, pBR-pLac-*gcvB*, pBR-pLac-*mcaS*, and pBR-pLac-*csrA*) was added to 100μL aliquots of cell suspension and incubated on ice for 30-minutes. Following the 30-minute incubation on ice, 900μL of LB was added to the cell-plasmid mixture, and the mixture was left to recover at 37°C for 1 hour on a shaking heat block. Upon the completion of the recovery period, 200μL of recovered cells were plated on LB-Amp plates and left to grow overnight at 37°C in a microbiological incubator. Transformants were purified once by streaking on LB agar plates supplemented with Ampicillin.

### RNA Isolation

Total RNA was isolated using the Hot Phenol Method as previously described (71, 72). Briefly, Overnight cultures were subcultured in 30 of LB at a 1:1000 dilution ratio and allowed to grow in the shaking water bath to an Optical Density of 600 (OD_600_) of 0.5. A 600ul aliquot of cells were isolated from exponentially growing *E. coli* cultures and mixed with resuspended in a 1X lysis buffer / Hot Acid Phenol solution in a 1.5 mL microcentrifuge tube on a thermomixer (Eppendorf) set to 65°C. The cells were incubated with intermittent shaking for 5 minutes. The tubes were subjected to centrifugation at 15,000 rpm for 10-minutes. The aqueous phase was extracted and purified two additional times with acid-phenol following by ethanol precipitation in −80°C freezer overnight. The RNA was pelleted and washed with 70% ethanol, air-dried, and resuspended with 50ul of DEPC water. RNA concentrations were successively measured using The NanoDrop^TM^ One^C^ Microvolume UV-Vis Spectrophotometer (Fisher Scientific).

### Agarose Northern Blot

The agarose based northern blot was executed as previously described (36, 73). A 1X MOPS (Quality Biological INC) 1% Agarose gel was used for the resolution of total RNA. After pre-running the gel at 100v for 40-minutes, a constant amount of total RNA (ranging from 2-5 μg) was mixed with 2X volume of loading buffer (500 μl Formamide, 100 μl 10X MOPS, 100 μl (80% glycerol 0.2% bromophenol blue), 120 μl Formaldehyde, 2 μl (10 mg/mL EtBr). The samples were then heated at 65° C for 15 minutes and loaded onto the gel for fractionation by gel electrophoresis for 40 minutes at 100 volts. The gel was then soaked in 0.05 M NaOH solution for 20 minutes and 20X SSC solution for 1-hour. The membrane was transferred to a Nylon Membrane using the capillary method as previously described (74). The membrane was then subjected to fixed to the membrane using the HL-2000 Hybrilinker (UVP). The crosslinked membrane was then pre-hybridized with 5 of ultrahyb oligo buffer (Ambion) for 2 hours, and then hybridized at 42°C in the HL-2000 Hybrilinker (UVP), after the addition of the transcript specific biotinylated oligonucleotide DNA probe. The membranes were then processed using stringency washes and developed using the Chemiluminescent Nucleic Acid Detection Module Kit (ThermoFisher Scientific) according to the manufacturer’s recommendations.

### β-galactosidase assays (Kinetic Microtiter Assays)

We measured β-galactosidase activity using the Kinetic Microtiter Plate Assay method (75). Briefly, overnight cultures were grown in 5 of LB Lennox liquid media at 37°C within a Cel-Gro Tissue culture rotator (Thermo Scientific) placed within a microbiological incubator (ShelLab). Overnight cultures containing sRNA plasmids were grown in LB-Amp. The following day, the overnight cultures were diluted 1:1000 in 30 mL of fresh LB or LB-Amp liquid media and placed in a 125 mL beveled Erlenmeyer flask to sub-culture the cells. IPTG was added to cultures containing sRNA plasmids to a final concentration of 1 mM. Cultures were then grown in a WS27 shaking water bath (ShelLab) at 37°C to an OD_600_ of 0.5. Cells were then harvested by centrifugation and then washed and resuspended with Lennox Broth (LB). The LB-cell suspensions were then transferred to a new flask and supplemented with ampicillin (100 μg/mL), IPTG (1 mM), and arabinose (0.02%). The cultures were then incubated in the 37°C shaking water bath for an additional 70 minutes. 100 μL aliquots of the cultures were collected once, or at 10-minute intervals, and transferred to a 96-well polystyrene microtiter plate containing 50μL of permeabilization solution (100 mM of Tris, pH 7.8, 32 mM of NaPO4, 8 mM of DTT, 8 mM CDTA, 4% Tris and 50 μL of Polymixin B). To serve as an experimental control, 100μL of LB was pipetted into one of the wells of the 96-well polystyrene microtiter plate. The 96-well polystyrene microtiter plate containing the samples was incubated at room temperature for 15-minutes to allow for cell lysis. Subsequently, 50μL of O-nitrophenyl-β-D-galactoside (ONPG) solution (4 mg/mL ONPG, 2 mM of Sodium Citrate, and 70 μL of β-Mercaptoethanol) was added to each well containing cell lysates. The 96-well polystyrene microtiter plate was immediately read using a Filter Max F5 Multi-Mode Micro Plate Reader (Molecular Devices).

### Statistical Analysis

All statistical analysis was executed using Prism 9 software (Graphpad).

## RESULTS

### CsrB was selected as a MiaA repressor in a targeted screen of small RNA regulators

We constructed a chromosomal *miaA*-*lacZ* translational fusion, whose expression is driven by the arabinose responsive P*_BAD_*promoter (Figure 1A). It was used for screening a plasmid-based small RNA library containing 30 *Escherichia coli* small RNAs that were cloned downstream of an IPTG inducible promoter (Mandin and Gottesman, 2009). The screening was carried out on Mac-Lac-Amp plates supplemented with arabinose to a final concentration of 0.002% to induce basal transcription from the P*_BAD_* promoter without producing a strong Lac^+^ phenotype, as previously described (76). This facilitated the identification of sRNA repressors more easily through our genetic screen. We identified five candidate sRNA regulators of MiaA: SdsR, ArcZ, GcvB, Spot42, and CsrB, with CsrB showing the strongest inhibitory effect (Figure 1B). To further test our hypothesis that post-transcriptional regulation of MiaA is modulated by one or more small RNAs, we executed quantiitative β-galactosidase assays of the P*_BAD_*-*miaA*-*lacZ* translational fusion strains carrying plasmids expressing SdsR, ArcZ, GcvB, Spot42 or CsrB (Figure 1C). Over-expression of ArcZ, GcvB, Spot42 did not affect P*_BAD_*-*miaA*-*lacZ* activity as compared to the vector control (Figure 1C). There was a 2-fold decrease in the activity of P*_BAD_*-*miaA*-*lacZ* translational fusion activity upon over-expression of CsrB or SdsR (Figure 1C), suggesting that CsrB and SdsR sRNAs are involved in the post-transcriptional repression of *miaA* expression. We also tested the impact of deleting the sRNA candidates on the expression of the P*_BAD_*-*miaA*-*lacZ* translational fusion (Figure 1D). Individual deletions of *gcvB*, *spf*, *arcZ*, or *sdsR* had no effect on the expression of the activity of the P*_BAD_*-*miaA*-*lacZ* translational fusion following arabinose induction (Figure 1D). The *csrB* mutant demonstated a modest increase in activity at 40 and 50 minutes after arabinose induction (Figures 1D and 2A). Given this modest increase in activity in the absence of *csrB*, we decided to focus our efforts on characterizing the CsrB effect on *miaA* expression, even though SdrS repressed activity of the fusion.

**Figure 1.**
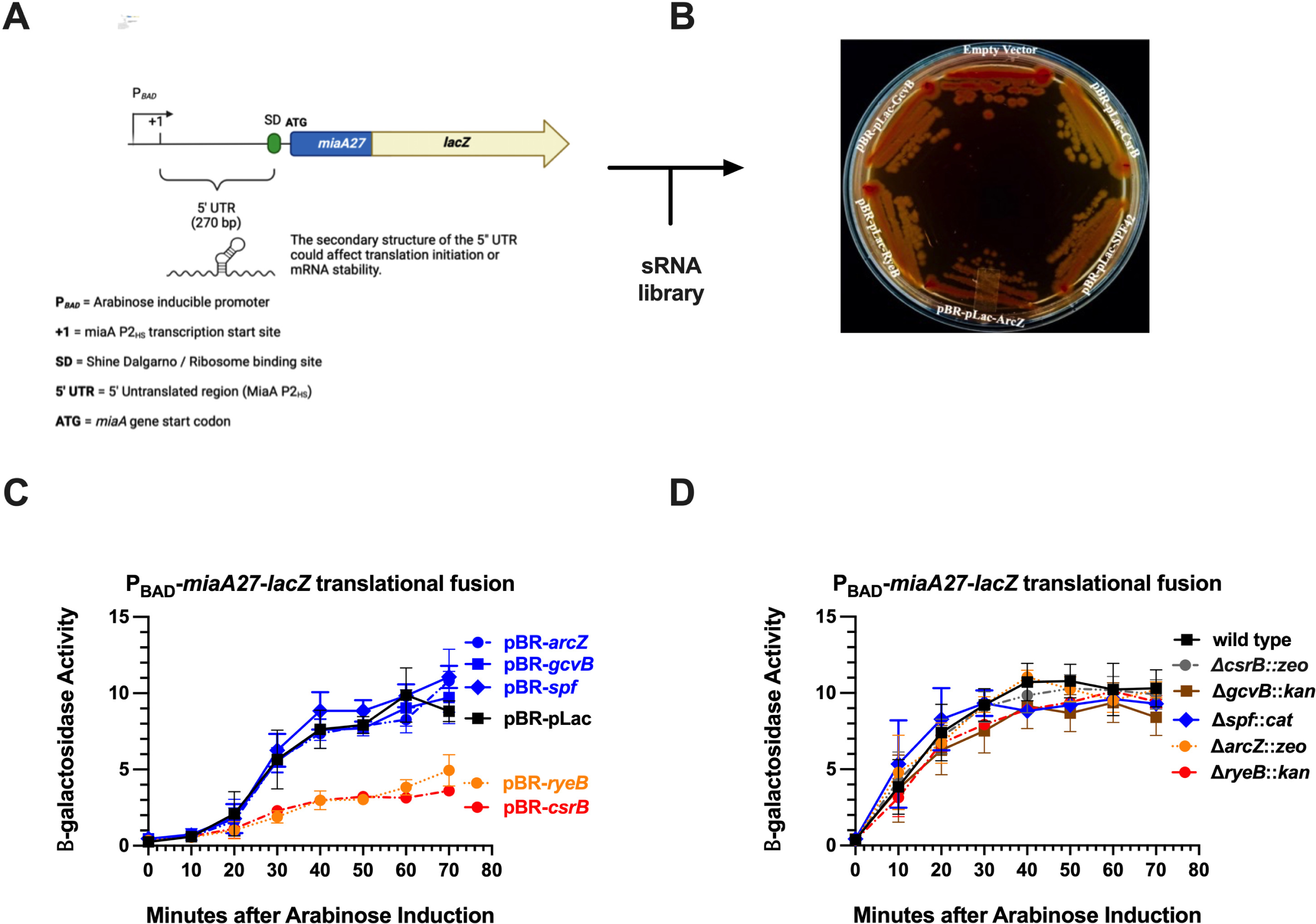
Small RNA Library Screen for Regulators of MiaA expression. A.) A schematic representing the arabinose-inducible *miaA*-*lacZ* translational gene fusion (P*_BAD_*-*miaA27*(_P2HS_)-*lacZ*) containing the 5’ UTR from the *miaA* P2 heat shock promoter B.) A P*_BAD_*-*miaA27*(_P2HS_)-*lacZ* translational fusion strain was transformed with a library of 30 known sRNAs cloned downstream from an IPTG-inducible promoter in plasmid pBR-pLac, and screened for activity on MacConkey-Lactose plates supplemented with Ampicillin. Results shown are for sRNA clones that gave a Lac-phenotype, suggesting a role for these small RNAs in the negative regulation of MiaA expression. C.) Quantitative β-galactosidase assay analysis of P*_BAD_*-*miaA27*(_P2HS_)-*lacZ* translational fusion with carrying pBR-*sdsR*, pBR-*arcZ*, pBR-*gcvB*, pBR-*spf*, or pBR-*csrB*. D.) Quantitative β-galactosidase assay analysis of P*_BAD_*-*miaA27*(_P2HS_)-*lacZ* translational fusion with deletion insertions in the genes for the candidate sRNA repressors of *miaA* picked up in our screen (Δ*csrB*::*zeo*, Δ*gcvB*::*kan*, Δ*spf*::*cat*, Δ*arcZ*::*zeo*, and Δ*sdsR*::*kan*).

**Figure 2.**
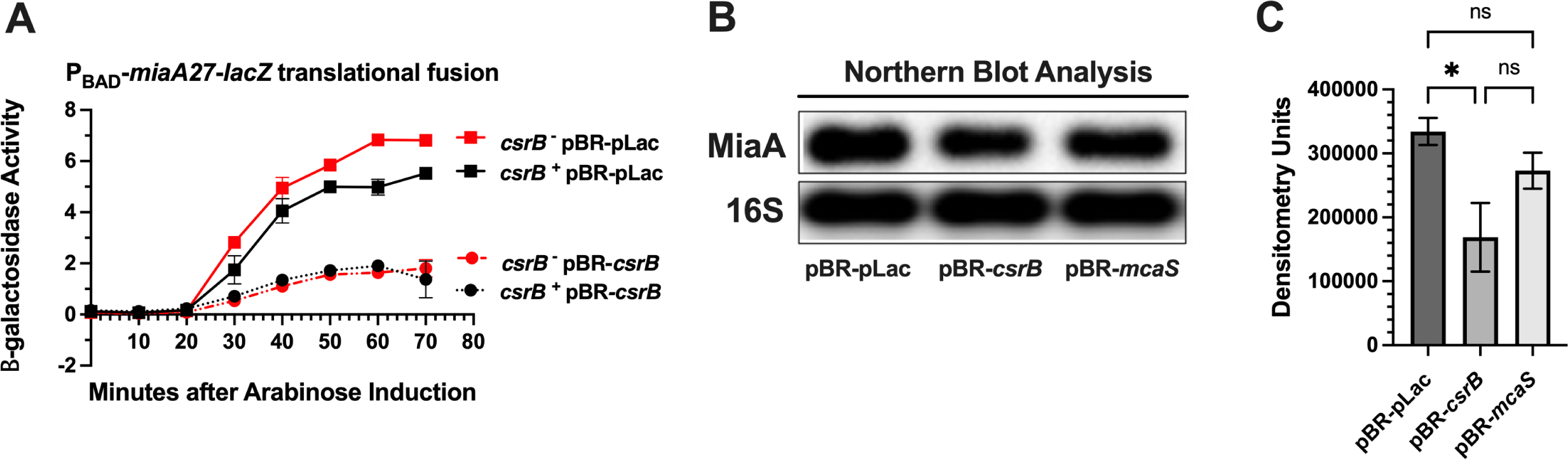
The effect of CsrB on MiaA expression. A.) Quantitative β-galactosidase assay analysis of P*_BAD_*-*miaA27*(_P2HS_)-*lacZ* translational fusion activity showing repression by CsrB sRNA. β-galactosidase assays were repeated at least three times and data points represent the mean + / -the standard error of the mean (mean +/-sem) B.) Northern blot analysis of MiaA steady state levels following over-expression of CsrB or McaS. Northern blots were repeated at least three times. C.) Quantitative densitometry of Northern blot analysis. Densitometry signals were acquired using a Fluorochem R Fluorescent / Chemiluminescent imager and statistical analysis of densitometry was executed using One Way ANOVA with Tukey’s Multiple Comparisons test on GraphPad Prism 9.

### CsrB affects *miaA* mRNA levels and translation

We transformed the the Δ*csrB* P*_BAD_*-*miaA*-*lacZ* translational fusion strain with pBR-pLac or pBR-*csrB* in order to execute and complementation assay (Figure 2A). We did observe complementation of the *csrB* mutant with the *csrB* plasmid. CsrB repression of *miaA*-*lacZ* was slightly more pronounced in the *csrB*^−^ background, causing a 7-fold vs 5-fold inhibition (Figure 2A). We then measured the steady-state levels of MiaA mRNA upon over-expression of CsrB and McaS. We decided to measure the McaS effect on MiaA mRNA levels since it also acts to sequester CsrA (77) (Figure 2B). Consistent with decreased activity of the P*_BAD_*-*miaA*-*lacZ* translational fusion, we observed a decrease of MiaA transcript level upon over-expression of CsrB (Figures 2B and 2C), with CsrB a much more effective regulator of the *miaA* RNA level. However, over-expression of McaS did not affect MiaA mRNA levels (Figures 2B and 2C).

### MiaA is regulated at the level of mRNA stability by PNPase and RNaseE

Since very little is known regarding the regulation of MiaA mRNA stability, we decided to test the role of RNsaes in MiaA mRNA turnover. Previous studies demonstrated an increase in mRNA levels of the *miaA* operon under non-permissive conditions in a temperature sensitive RNaseE mutant (31). We tested the roles of RNaseE and PNPase, both of which are components of the RNA Degradosome, in *miaA* mRNA turnover. Specifically, we compared *miaA* mRNA recycling in the cell, in WT isogenic wild type, PNPase mutant (*pnpA*^-^), and temperature sensitive RnaseE mutants (*rne^ts^*) (Figure 3). The wild type and *rne^ts^* strains were grown at 32°C to mid-log phase and then shifted to 43.5°C, the non-permissive condition for the RnaseE *ts* mutant. Then, we immediately added rifampicin to the cultures to halt transcription. We then isolated total RNA at times of 0, 2, 4, 8, 16, and 32 minutes after rifampicin treatment and measured MiaA mRNA levels by northern blot (Figure 3A and 3B). Upon semi-quantitative densitometric analysis of the Northern Blot analysis and statistical analysis of the MiaA mRNA in this experiment (Figures 3B and 3C), we determined that the *t*_1/2_ of the MiaA mRNA increased in the *rne*^ts^ (> 32 minutes) vs wild type control (17 minutes) (Figure 3C). We grew wild type and Δ*pnpA*::*kan* mutants in rich media at 37°C to mid-log, added rifampicin, and isolated total RNA at 0, 2, 4, 8, 16, and 32 minutes post rifampicin treatment. The *t*_1/2_ of the MiaA mRNA is > 32 minutes and 20 minutes, in the Δ*pnpA*::*kan* and wild type genetic backgrounds, respectively (Figure 3B). The longer half-life of the MiaA transcript in the *rne*^ts^ allele and Δ*pnpA*::*kan* mutant, in comparison to the wild type control, suggests that RNaseE and PNPase are involved in the post-transcriptional regulation of MiaA at the level of mRNA stability.

**Figure 3.**
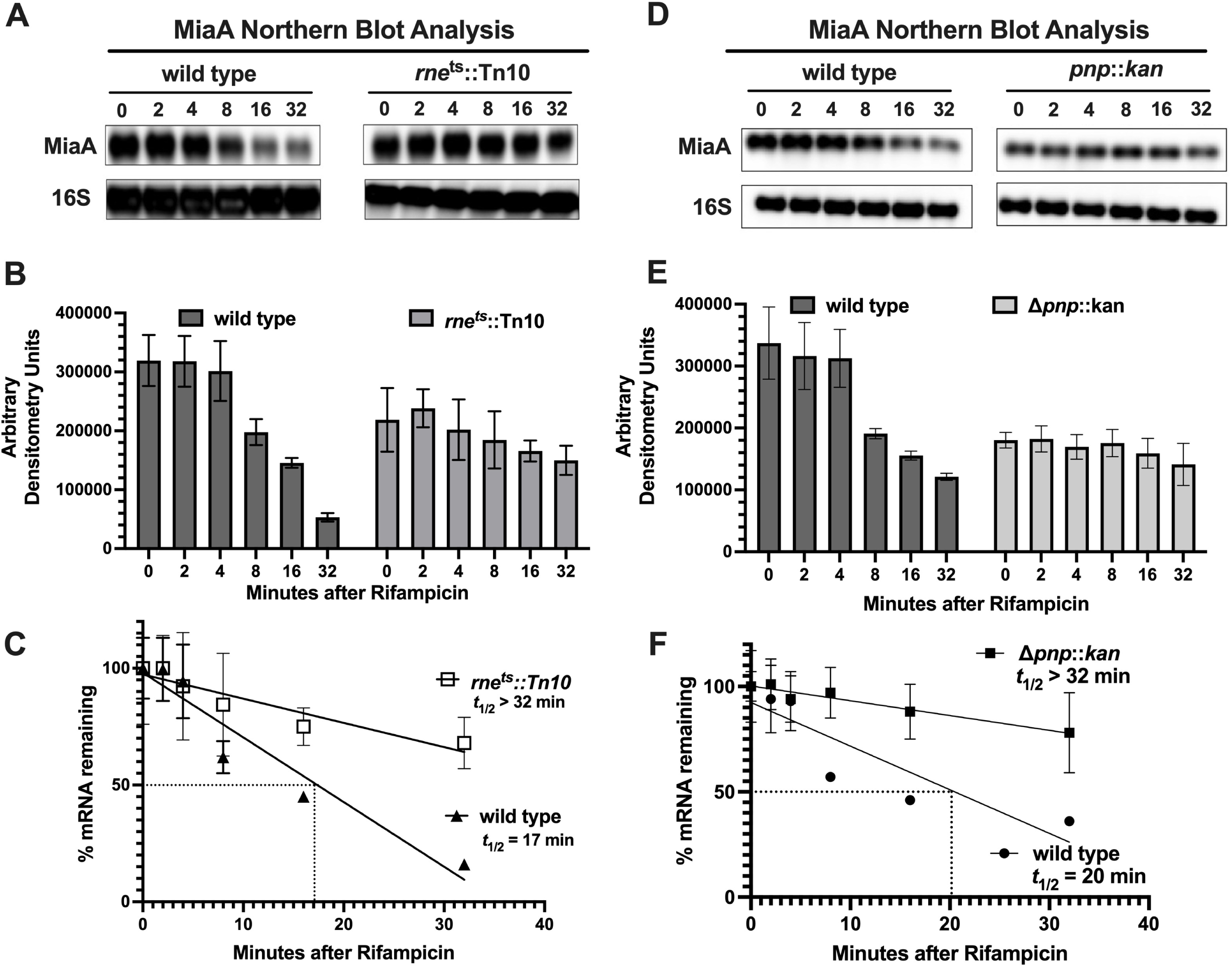
Effects of RNaseE and PNPase on MiaA mRNA stability. A) Northern Blot analysis of MiaA mRNA stability. A *rne-3071 zce-726*::Tn*10* allele was transduced into MG1655 by bacteriophage P1 transduction and selected for tetracycline (Tet^R^) resistance. Strains were grown in rich media (LB) at 30°C to an OD_600_ of 0.3. 600ul sample aliquots were collected for total RNA isolation and northern blot analysis at zero-minutes, before transferring cultures to 43°C. Rifampicin was added, and samples were collected at 2,4,8,16, and 32 minutes for total RNA isolation and analysis by Agarose Northern Blot. Experiments were repeated at least three times and blots shown are a representative blot of the triplicate. B) Quantitative densitometry of Northern Blot in section A. Statistical analysis includes mean and standard error of the mean (mean + / -s.e.m.) C.) Half-life calculations of Northern Blot executed in Section A. Quantitative Northern Blot data from section B was subjected to linear regression analysis using GraphPad Prism 9. D.) Δ*pnp*::*kan* mutation was transduced into MG1655 by bacteriophage P1 transduction and selected for kanamycin (kan^R^) resistance. Strains were grown in rich media (LB) to an OD_600_ of 0.3. 600ul aliquots of the sample were collected for total RNA isolation and northern blot analysis at zero-minutes. Rifampicin was added, and samples were collected at 2,4,8, and 16 minutes for total RNA isolation and analysis by Agarose Northern Blot. E) Quantitative densitometry of Northern Blot in section A. Statistical analysis includes mean and standard error of the mean (mean + / -s.e.m.) F.) Half-life calculations of Northern Blot executed in Section D. Quantitative Northern Blot data from Section B were subjected to linear regression analysis using GraphPad Prism 9.

### CsrA is necessary for the full expression of MiaA

The RNA binding protein CsrA is part of a global regulatory system that controls bacterial gene expression at the post-transcriptional level (reference). CsrA regulates translation of target proteins by binding to target sequences in the 5’ UTR of the target genes. The availability of CsrA is regulated by binding of to CsrB and CsrC sRNAs. CsrB and CsrC have multiple CsrA binding sites, each binding to approximately 18 CsrA subunits and inhibiting the activity of CsrA. Since CsrB represses the expression of MiaA, and CsrB acts to sequester and inhibit CsrA activity, we hypothesized that the CsrB regulatory effect on MiaA may be through CsrA. To test our hypothesis, we measured the activity of the P*_BAD_*-*miaA27*-*lacZ* translational fusion in a *csrA*^-^genetic background over-expression of CsrA (Figure 4). In the *csrA*^-^background, the β-galactosidase activity of the P*_BAD_*-*miaA27*-*lacZ* strain was decreased by an approximate 6-10-fold decrease in β-galactosidase activity after 70 minutes after arabinose induction, and was essentially non-detectable, in comparison to the *csrA*^+^ and *csrB*^-^genetic backgrounds (Figure 4A). The β-galactosidase activity of the *csrA*^-^P*_BAD_*-*miaA27*-*lacZ* strain was partially rescued by over-expression of plasmid-based *csrA* (Figure 4B). Given the decrease in the activity of the P*_BAD_*-*miaA27*-*lacZ* fusion, in the *csrA* mutant, we decided to measure miaA mRNA levels the absence *csrA*. We measured *miaA* mRNA levels in the wild type, *csrA*^−^, and *csrB*^−^ genetic backgrounds. The *miaA* mRNA levels were decreased, in a statistically significant manner, by approximately 20-fold in the absence of *csrA* while remaining virtually unchanged in the absence of *csrB* (Figures 4C and 4D). This result is consistent with MiaA mRNA down-regulation upon over-expression of CsrB (Figure 1C) and confirms that the CsrA-CsrB system regulates post-transcriptional expression of MiaA.

**Figure 4.**
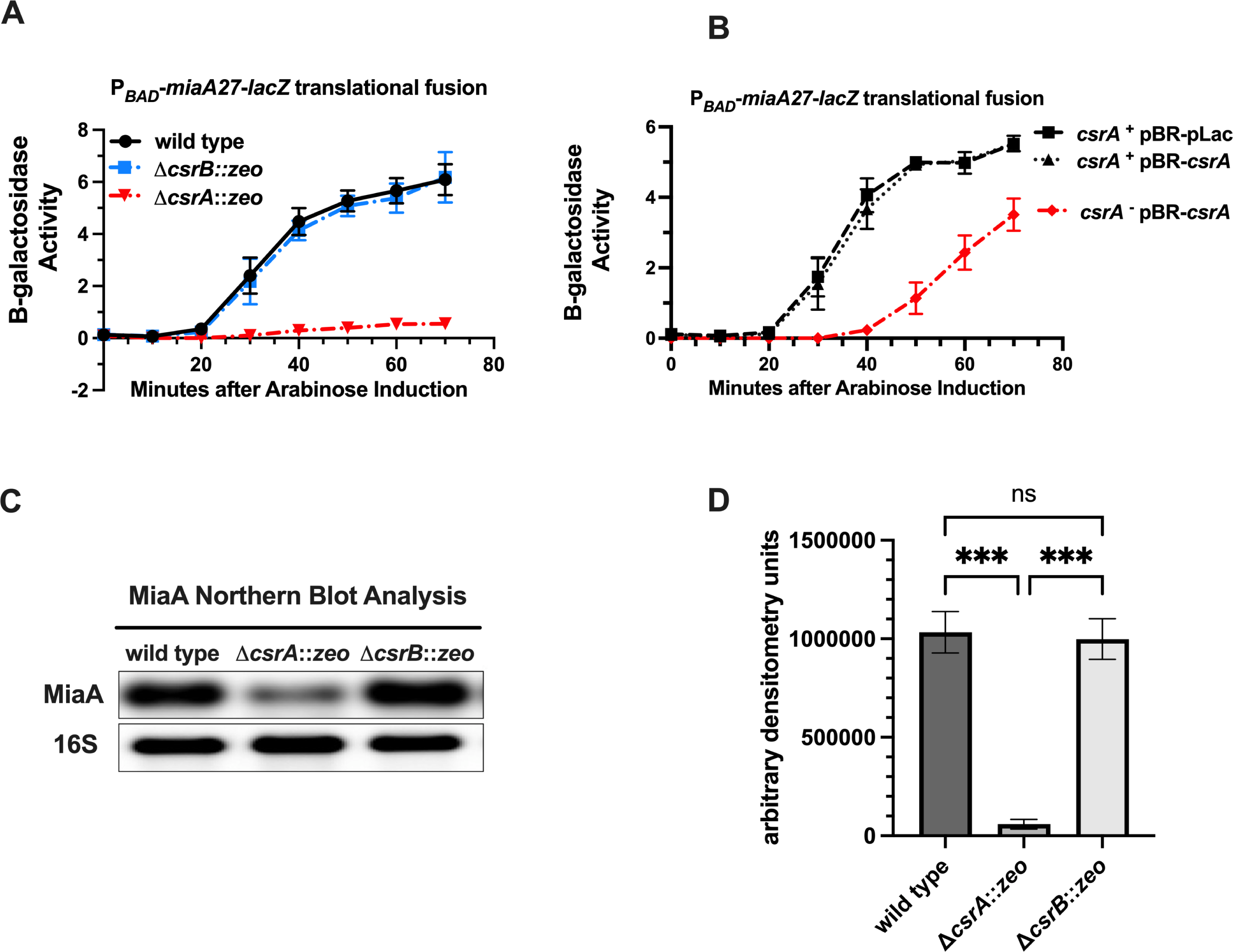
MiaA Expression in the absence of *csrA*. (A) Wild type, Δ*csrA,* and Δ*csrB* versions of P*_BAD_*-*miaA27*(_P2HS_)-*lacZ* translational fusions were grown in LB supplemented with glucose to an OD_600_ of 0.5, shifted to LB supplemented with arabinose and aliquots were obtained for β-galactosidase assay every 10 minutes for 70 minutes. (B) CsrA complementation assay using P*_BAD_*-*miaA27*(_P2HS_)-*lacZ* translational fusion activity. Wild type and Δ*csrA*::*zeo* strains of P*_BAD_*-*miaA27*(_P2HS_)-*lacZ* fusions carrying pBR-pLac or pBR-*csrA* were grown in rich media (LB) to an OD_600_ of 0.5 and shifted to LB supplemented with arabinose and aliquots were obtained for β-galactosidase assay every 10 minutes for 70 minutes. (C) Northern Blot analysis of MiaA from total RNA isolated from exponentially growing wild type, Δ*csrA,* and Δ*csrB* cells. Experiments were repeated at least three times and blots shown are a representative blot of the triplicate. (D.) Quantitative densitometry of Northern blot analysis in (C). Densitometry signals were acquired using Fluorochem R Fluorescent / Chemiluminescent imager and statistical analysis of densitometry was executed using One Way ANOVA with Tukey’s Multiple Comparisons test on GraphPad Prism 9.

## DISCUSSION

### RNA modifications and The Bacterial Epitranscriptome

RNA modifications have been classically demonstrated to promote translational fidelity and the stability of tRNAs and rRNAs. Yet, contemporary studies have uncovered an expanded functional scope for RNA modifications. Studies in eukaryotic systems have highlighted an expanded list of RNA species that contain RNA modifications. Specifically, RNA modifications have been found within eukaryotic regulatory RNA species such as miRNA, CircRNAs, and lncRNAs (78–80). They are thought to play roles in post-transcriptional regulatory circuits by modulating miRNA interactions with the 3’ UTRs of mRNAs, modulating mRNA stability. Several different RNA modifications play regulatory roles in cellular physiology (81–86). Work by our group and others has demonstrated that RNA modifications may play regulatory roles in gene expression by promoting the expression of stress response genes, in a manner dependent upon codon bias in bacteria and bacteriophage (25–27, 87–89). Subsequent reports from several groups have also demonstrated codon biased gene regulation in bacterial pathogens (25–27, 89, 90). In all of these studies, the role of the modification was established by mutating the gene(s) encoding the modification enzymes. To add to these discoveries, understanding the conditions whereby RNA modifications are synthesized will assist us in understanding their impact on the physiology of the cell.

### New regulators of MiaA expression

We previously demonstrated that MiaA is necessary for the expression of RpoS, IraP, and Hfq (25–27). The MiaA requirement for optimal expression of RpoS and IraP is related UUX leucine decoding, suggesting that MiaA may promote the expression of stress response genes during leucine starvation (25–27). Over-expression of leucine tRNAs were able to suppress the decreased expression of *rpoS* in *miaA* mutants. Interestingly, a previous study demonstrated that *miaA* and *leuX* (leucine tRNA) are synthetically lethal during heat shock (91). Also, *miaA* mutants affect leucine operon gene expression in Salmonella (4). Taken together, this suggests that MiaA catalyzed i^6^A37 modification is particularly important during heat shock whereby UUX leucine decoding provides an adaptive advantage. For these reasons, characterizing the post-transcriptional regulation of the MiaA transcript from the σ^32^ dependent heat shock promoter is of critical importance. Prior to this work, little was known about post-transcriptional regulation of MiaA. Here, we started to fill that knowledge gap by identifying several potential candidate small RNA regulators of MiaA expression at the post-transcriptional level. Our results demonstrate a strong effect of CsrA, likely as a direct post-transcriptional regulator of the *miaA*_P2_ promoter transcript.

### MiaA as an additional potential stimulatory target of the CsrA-CsrB system and interactions with other post-transriptional regulators of the *miaA* operon

CsrA can directly bind to mRNA transcripts to regulate gene expression at the post-transcriptional level, in the absence of sequestration by CsrB or CsrC (40). There are approximately 12 mRNA transcripts that are post-transcriptionally regulated by CsrA. 10 of these regulatory targets are repressed and two of these regulatory targets are stimulated by CsrA, including *ymdA* (92) (Table 3). CsrA stimulates the translation of *ymdA* thorugh interaction with its 5’ UTR to reverse the formation of secondary structures that occlude its ribosome binding site and subsequent translational initiation. Our work identifies MiaA as an additional stimulatory target of CsrA. It is possible that the 5’ UTR of the *miaA*_P2_ transcript has secondary structures that occlude the ribosome binding site and that CsrA acts in a similar manner on this transcript to stimulate translation of this transcript. Given the fact that RNaseE and PNPase mutants result in the stabilization of the MiaA transcript, CsrA may also interact with Degradosome enzymes or the 5’ untranslated region (UTR) of the MiaA P2 transcript to promote its stabilization. We have illustrated a simple model to describe CsrA-CsrB regulation of MiaA (Figure 5). Upon sequestration of CsrA by CsrB, *miaA* transcript from the P2 (heat shock promoter) is less stable and translation is also inhibited. This is reversed in the absence of CsrB, and when CsrA levels are higher or its more active, resulting in transcript stabilization and an increase in translation (Figure 5). The stabilization of the miaA mRNA could be through antagonistic interactions with Degradosome proteins RNaseE and PNPase. This model is consistent with the previous report from Vakulskas et. al. (2016), whereby CsrA was shown to stabilize CsrB through antagonistic interactions with RNaseE (93).

**Figure 5.**
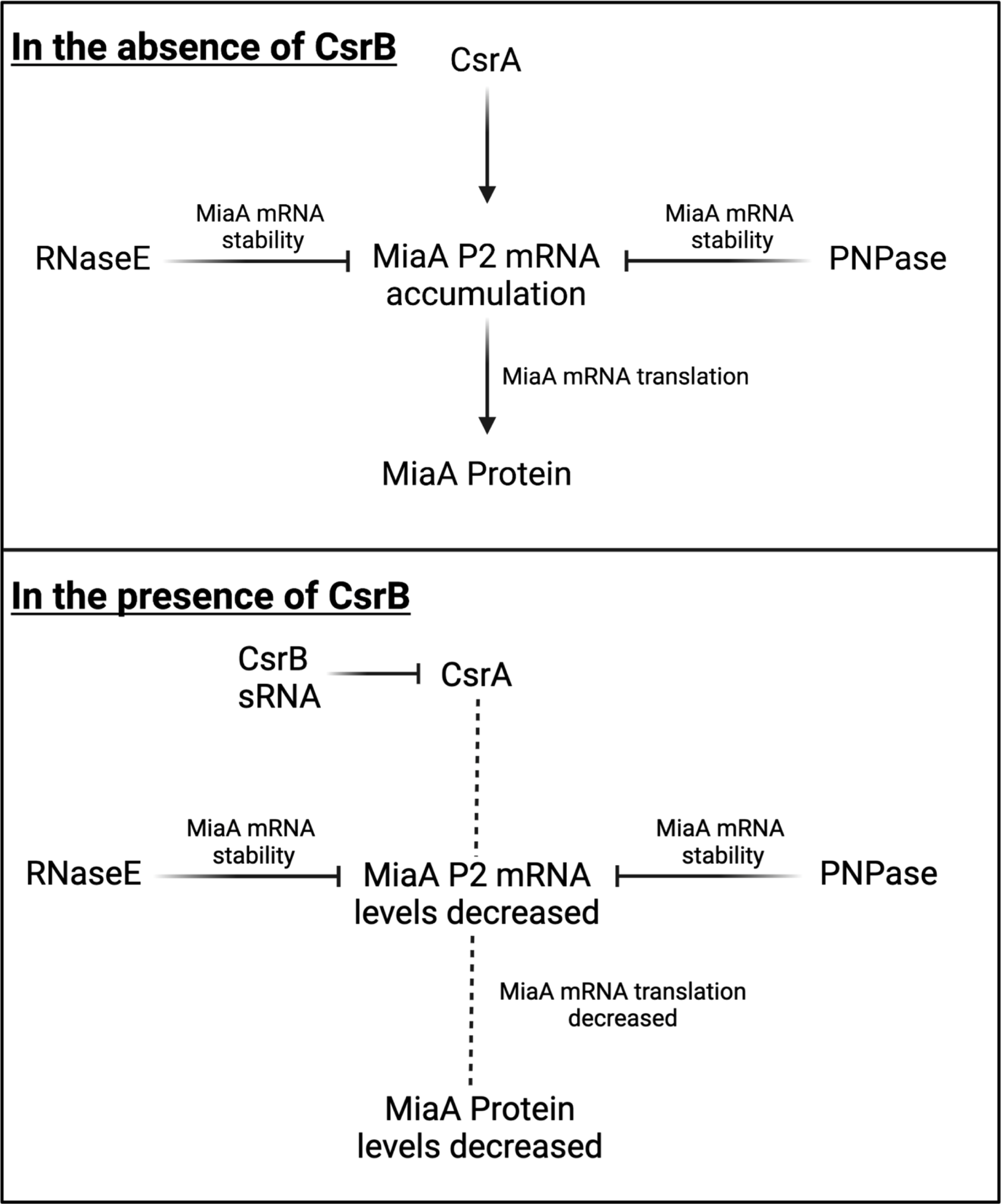
Model for CsrA-CsrB regulation of MiaA expression. Graphical model of CsrA / CsrB regulation of MiaA was created using Biorender. MiaA transcript turnover is mediated through both PNPase and RNaseE. CsrA promotes the accumulation of the MiaA P2 (Heat Shock) (HS) mRNA in the absence of CsrB. In the presence of CsrB, CsrA sequestration leads to decreased levels of the MiaA P2 (Heat Shock) mRNA.

**Table 3.-.** CsrA regulated.

| **<u>Repressed</u>** | **<u>Activated</u>** |
| --- | --- |
| *glgC* | *moaABC operon* |
| *glgS* | *ymdAB-clsC* |
| *sdiA* |  |
| *dgcZ* |  |
| *dgcT* |  |
| *pgaA* |  |
| *pdeI* |  |
| *nhaR* |  |
| *rpoE* |  |
| *iraD* |  |

The *miaA* gene is encoded directly upstream of the *hfq* gene within the *nnr*-*tsaE*-*amiB*-*mutL*-*miaA*-*hfq*-*hflX*-*hflK*-*hflC* superoperon in *E. coli* genome (30, 31, 94, 95). This tandem genetic localization is widely conserved across the the prokaryotic domain. While mutations in *hfq* exhibit extensively pleiotropic effects, *miaA* mutant phenotypes appear to be distinct with the exception to its effects on *rpoS* expression whereby the *miaA* mutant demonstrates some polarity likely due to effects on Hfq-dependent sRNAs that stimulation *rpoS* translation (25, 96–101). There is complex transcriptional organization of this superoperon with approximately 10 promoters exhibiting alternating dependence upon σ^32^ an σ^70^, between the *amiB*-*hfq* region (30, 31, 94, 95). All of these promoters are upstream of the *hfq* gene. However, there are three promoters between the *miaA* gene and the start codon of *hfq*, with the function of drive expression of *hfq*-*hflXKC*: defined as *hfq*_P1(hs)_, *hfq*_P3_, *hfq*_P3_. Baker et. al. (2007) demonstrating that CsrA binds to the ribosome binding site of the *hfq* mRNA and inhibits *hfq* translation (102). In our screen, we utilized a *miaA*-*lacZ* fusion with the *miaA* P2 transcript. If CsrA interacts with the 5’ UTR of the *miaA* P2 transcript, it may provide CsrA with multiple options for binding to the larger transcripts expressed from this operon that contains both *miaA* and *hfq*. Multiple CsrA binding sites may or may not be accessible on the larger trancripts, depending on their. This is the subject of ongoing studies in our laboratory.

## Acknowledgements

We would like to thank the members of the Thompson Laboratory, Muneer Abbas, Qiyi Tang, Nadim Majdalani, and Susan Gottesman for critical review of this manuscript.

## Funding

This work was supported by Award Number 1904089 from the National Science Foundation (NSF).

## References

1. Caillet J, Droogmans L. 1988. Molecular cloning of the *Escherichia coli miaA* gene involved in the formation of delta 2-isopentyl adenosine in tRNA. J Bacteriol 170:4147–4152.

2. Connolly DM, Winkler ME. 1989. Genetic and physiological relationships among the *miaA* gene, 2-methylthio-N6-(delta 2-isopentenyl)-adenosine tRNA modification, and spontaneous mutagenesis in *Escherichia coli* K-12. J Bacteriol 171:3233–46.

3. Connolly DM, Winkler ME. 1991. Structure of *Escherichia coli* K-12 *miaA* and characterization of the mutator phenotype caused by *miaA* insertion mutations. J Bacteriol 173:1711–1721.

4. Blum PH. 1988. Reduced *leu* operon expression in a *miaA* mutant of *Salmonella typhimurium*. J Bacteriol 170:5125–5133.

5. Diaz I, Ehrenberg M, Kurland CG. 1986. How do combinations of rpsL- and miaA-generate streptomycin dependence? Mol Gen Genet 202:207–11.

6. Pierrel F, Bjork GR, Fontecave M, Atta M. 2002. Enzymatic modification of tRNAs: MiaB is an iron-sulfur protein. J Biol Chem 277:13367–70.

7. Esberg B, Leung HC, Tsui HC, Bjork GR, Winkler ME. 1999. Identification of the *miaB* gene, involved in methylthiolation of isopentenylated A37 derivatives in the tRNA of *Salmonella typhimurium* and *Escherichia coli*. J Bacteriol 181:7256–65.

8. Tukenmez H, Xu H, Esberg A, Bystrom AS. 2015. The role of wobble uridine modifications in +1 translational frameshifting in eukaryotes. Nucleic Acids Res 43:9489–9499.

9. Bjork GR, Durand JM, Hagervall TG, Leipuviene R, Lundgren HK, Nilsson K, Chen P, Qian Q, Urbonavicius J. 1999. Transfer RNA modification: influence on translational frameshifting and metabolism. FEBS Lett 452:47–51.

10. Bjork GR, Wikstrom PM, Bystrom AS. 1989. Prevention of translational frameshifting by the modified nucleoside 1-methylguanosine. Science 244:986–9.

11. Urbonavicius J, Qian Q, Durand JM, Hagervall TG, Bjork GR. 2001. Improvement of reading frame maintenance is a common function for several tRNA modifications. EMBO J 20:4863–4873.

12. Urbonavicius J, Stahl G, Durand JM, Ben Salem SN, Qian Q, Farabaugh P, Bjork GR. 2003. Transfer RNA modifications that alter +1 frameshifting in general fail to affect −1 frameshifting. RNA 9:760–768.

13. Qian Q, Bjork GR. 1997. Structural requirements for the formation of 1-methylguanosine in vivo in tRNA(Pro)GGG of Salmonella typhimurium. J Mol Biol 266:283–96.

14. Qian Q, Bjork GR. 1997. Structural alterations far from the anticodon of the tRNAProGGG of *Salmonella typhimurium* induce +1 frameshifting at the peptidyl-site. J Mol Biol 273:978–92.

15. Qian Q, Curran JF, Bjork GR. 1998. The methyl group of the N6-methyl-N6-threonylcarbamoyladenosine in tRNA of *Escherichia coli* modestly improves the efficiency of the tRNA. J Bacteriol 180:1808–13.

16. Zhao J, Leung HE, Winkler ME. 2001. The *miaA* mutator phenotype of *Escherichia coli* K-12 requires recombination functions. J Bacteriol 183:1796–800.

17. Schweizer U, Bohleber S, Fradejas-Villar N. 2017. The modified base isopentenyladenosine and its derivatives in tRNA. RNA Biol 14:1197–1208.

18. Nishii K, Wright F, Chen YY, Moller M. 2018. Tangled history of a multigene family: The evolution of ISOPENTENYLTRANSFERASE genes. PLoS One 13:e0201198.

19. Soman S, Ram S. 2022. MiaA (Rv2727c) mediated tRNA isopentenylation of *Mycobacterium tuberculosis* H37Rv. Mol Biol Res Commun 11:97–104.

20. Fleming BA, Blango MG, Rousek AA, Kincannon WM, Tran A, Lewis AJ, Russell CW, Zhou Q, Baird LM, Barber AE, Brannon JR, Beebout CJ, Bandarian V, Hadjifrangiskou M, Howard MT, Mulvey MA. 2022. A tRNA modifying enzyme as a tunable regulatory nexus for bacterial stress responses and virulence. Nucleic Acids Res 50:7570–7590.

21. Durand JM, Bjork GR, Kuwae A, Yoshikawa M, Sasakawa C. 1997. The modified nucleoside 2-methylthio-N6-isopentenyladenosine in tRNA of *Shigella flexneri* is required for expression of virulence genes. J Bacteriol 179:5777–5782.

22. Sun B, Liu H, Jiang Y, Shao L, Yang S, Chen D. 2020. New Mutations Involved in Colistin Resistance in *Acinetobacter baumannii*. mSphere 5:e00895–19.

23. Koshla O, Yushchuk O, Ostash I, Dacyuk Y, Myronovskyi M, Jager G, Sussmuth RD, Luzhetskyy A, Bystrom A, Kirsebom LA, Ostash B. 2019. Gene *miaA* for post-transcriptional modification of tRNA(XXA) is important for morphological and metabolic differentiation in *Streptomyces*. Mol Microbiol 112:249–265.

24. Koshla O, Kravets V, Dacyuk Y, Ostash I, Sussmuth R, Ostash B. 2020. Genetic analysis of *Streptomyces albus* J1074 *mia* mutants suggests complex relationships between post-transcriptional tRNA(XXA) modifications and physiological traits. Folia Microbiol (Praha) 65:1009–1015.

25. Thompson KM, Gottesman S. 2014. The MiaA tRNA modification enzyme is necessary for robust RpoS expression in *Escherichia coli*. J Bacteriol 196:754–61.

26. Aubee JI, Olu M, Thompson KM. 2017. TrmL and TusA are necessary for rpoS and MiaA is required for hfq expression in *Escherichia coli*. Biomolecules 7:39.

27. Aubee JI, Olu M, Thompson KM. 2016. The i^6^A37 tRNA modification is essential for proper decoding of UUX-Leucine codons during *rpoS* and *iraP* translation. RNA 22:729–742.

28. Moller T, Franch T, Hojrup P, Keene DR, Bachinger HP, Brennan RG, Valentin-Hansen P. 2002. Hfq: a bacterial Sm-like proteins that mediates RNA-RNA interaction. Mol Cell 9:23–30.

29. Kajitani M, Kato A, Wada A, Inokuchi H, Ishihama A. 1994. Regulation of the *Escherichia coli hfq* gene encoding the host factor for phage Q beta. J Bacteriol 176:531–534.

30. Tsui HC, Feng G, Winkler ME. 1996. Transcription of the *mutL* repair, *miaA* tRNA modification, *hfq* pleiotropic regulator, and *hflA* region protease genes of Escherichia coli K-12 from clustered Esigma32-specific promoters during heat shock. J Bacteriol 178:5719–31.

31. Tsui HC, Winkler ME. 1994. Transcriptional patterns of the *mutL-miaA* superoperon of *Escherichia coli* K-12 suggest a model for posttranscriptional regulation. Biochimie 76:1168–77.

32. Tsui HC, Feng G, Winkler ME. 1997. Negative regulation of *mutS* and *mutH* repair gene expression by the Hfq and RpoS global regulators of *Escherichia coli* K-12. J Bacteriol 179:7476–87.

33. Py B, Causton H, Mudd EA, Higgins CF. 1994. A protein complex mediating mRNA degradation in *Escherichia coli*. Mol Microbiol 14:717–29.

34. Py B, Higgins CF, Krisch HM, Carpousis AJ. 1996. A DEAD-box RNA helicase in the Escherichia coli RNA degradosome. Nature 381:169–72.

35. Mudd EA, Higgins CF. 1993. *Escherichia coli* endoribonuclease RNase E: autoregulation of expression and site-specific cleavage of mRNA. Mol Microbiol 9:557–68.

36. Masse E, Escorcia FE, Gottesman S. 2003. Coupled degradation of a small regulatory RNA and its mRNA targets in *Escherichia coli*. Genes Dev 17:2374–83.

37. Guillier M, Gottesman S, Storz G. 2006. Modulating the outer membrane with small RNAs. Genes Dev 20:2338–48.

38. Mandin P, Gottesman S. 2010. Integrating anaerobic/aerobic sensing and the general stress response through the ArcZ small RNA. EMBO J 29:3094–107.

39. Luo X, Majdalani N. 2024. Directed Screening for sRNA Targets in *E. coli* Using a Plasmid Library. Methods Mol Biol 2741:291–306.

40. Liu MY, Gui G, Wei B, Preston JF, 3rd, Oakford L, Yuksel U, Giedroc DP, Romeo T. 1997. The RNA molecule CsrB binds to the global regulatory protein CsrA and antagonizes its activity in *Escherichia coli*. J Biol Chem 272:17502–10.

41. Liu MY, Romeo T. 1997. The global regulator CsrA of *Escherichia coli* is a specific mRNA-binding protein. J Bacteriol 179:4639–42.

42. Sabnis NA, Yang H, Romeo T. 1995. Pleiotropic regulation of central carbohydrate metabolism in *Escherichia coli* via the gene *csrA*. J Biol Chem 270:29096–104.

43. Pannuri A, Vakulskas CA, Zere T, McGibbon LC, Edwards AN, Georgellis D, Babitzke P, Romeo T. 2016. Circuitry Linking the Catabolite Repression and Csr Global Regulatory Systems of *Escherichia coli*. J Bacteriol 198:3000–3015.

44. Wang X, Dubey AK, Suzuki K, Baker CS, Babitzke P, Romeo T. 2005. CsrA post-transcriptionally represses pgaABCD, responsible for synthesis of a biofilm polysaccharide adhesin of *Escherichia coli*. Mol Microbiol 56:1648–63.

45. Wei BL, Brun-Zinkernagel AM, Simecka JW, Pruss BM, Babitzke P, Romeo T. 2001. Positive regulation of motility and *flhDC* expression by the RNA-binding protein CsrA of *Escherichia coli*. Mol Microbiol 40:245–56.

46. Wang D, McAteer SP, Wawszczyk AB, Russell CD, Tahoun A, Elmi A, Cockroft SL, Tollervey D, Granneman S, Tree JJ, Gally DL. 2018. An RNA-dependent mechanism for transient expression of bacterial translocation filaments. Nucleic Acids Res 46:3366–3381.

47. Liaw SJ, Lai HC, Ho SW, Luh KT, Wang WB. 2003. Role of RsmA in the regulation of swarming motility and virulence factor expression in *Proteus mirabilis*. J Med Microbiol 52:19–28.

48. Nava-Galeana J, Yakhnin H, Babitzke P, Bustamante VH. 2023. CsrA Positively and Directly Regulates the Expression of the *pdu*, *pocR*, and *eut* Genes Required for the Luminal Replication of *Salmonella Typhimurium*. Microbiol Spectr 11:e0151623.

49. Zhu D, Wang S, Sun X. 2021. FliW and CsrA Govern Flagellin (FliC) Synthesis and Play Pleiotropic Roles in Virulence and Physiology of Clostridioides difficile R20291. Front Microbiol 12:735616.

50. Hubloher JJ, Schabacker K, Muller V, Averhoff B. 2021. CsrA Coordinates Compatible Solute Synthesis in *Acinetobacter baumannii* and Facilitates Growth in Human Urine. Microbiol Spectr 9:e0129621.

51. Butz HA, Mey AR, Ciosek AL, Crofts AA, Davies BW, Payne SM. 2021. Regulatory Effects of CsrA in *Vibrio cholerae*. mBio 12.

52. Dai Q, Xu L, Xiao L, Zhu K, Song Y, Li C, Zhu L, Shen X, Wang Y. 2018. RovM and CsrA Negatively Regulate Urease Expression in *Yersinia pseudotuberculosis*. Front Microbiol 9:348.

53. Potts AH, Leng Y, Babitzke P, Romeo T. 2018. Examination of Csr regulatory circuitry using epistasis analysis with RNA-seq (Epi-seq) confirms that CsrD affects gene expression via CsrA, CsrB and CsrC. Sci Rep 8:5373.

54. Heroven AK, Bohme K, Dersch P. 2012. The Csr/Rsm system of Yersinia and related pathogens: a post-transcriptional strategy for managing virulence. RNA Biol 9:379–91.

55. Abbott ZD, Yakhnin H, Babitzke P, Swanson MS. 2015. *csrR*, a Paralog and Direct Target of CsrA, Promotes Legionella pneumophila Resilience in Water. mBio 6:e00595.

56. Ozturk G, LeGrand K, Zheng Y, Young GM. 2017. Yersinia enterocolitica CsrA regulates expression of the Ysa and Ysc type 3 secretion system in unique ways. FEMS Microbiol Lett 364.

57. Muller P, Gimpel M, Wildenhain T, Brantl S. 2019. A new role for CsrA: promotion of complex formation between an sRNA and its mRNA target in *Bacillus subtilis*. RNA Biol 16:972–987.

58. Heroven AK, Nuss AM, Dersch P. 2017. RNA-based mechanisms of virulence control in Enterobacteriaceae. RNA Biol 14:471–487.

59. Vakulskas CA, Potts AH, Babitzke P, Ahmer BM, Romeo T. 2015. Regulation of bacterial virulence by Csr (Rsm) systems. Microbiol Mol Biol Rev 79:193–224.

60. Cerca N, Jefferson KK. 2008. Effect of growth conditions on poly-N-acetylglucosamine expression and biofilm formation in *Escherichia coli*. FEMS Microbiol Lett 283:36–41.

61. Lee JH, Ancona V, Chatnaparat T, Yang H, Zhao Y. 2019. The RNA-binding protein CsrA controls virulence in *Erwinia amylovora* by regulating RelA, RcsB, and FlhD at the posttranscriptional level. Mol Plant Microbe Interact 32:1448–1459.

62. Rojano-Nisimura AM, Simmons TR, Leistra AN, Mihailovic MK, Buchser R, Ekdahl AM, Joseph I, Curtis NC, Contreras LM. 2023. CsrA selectively modulates sRNA-mRNA regulator outcomes. Front Mol Biosci 10:1249528.

63. Stenum TS, Holmqvist E. 2022. CsrA enters Hfq’s territory: Regulation of a base-pairing small RNA. Mol Microbiol 117:4–9.

64. Lai YJ, Yakhnin H, Pannuri A, Pourciau C, Babitzke P, Romeo T. 2022. CsrA regulation via binding to the base-pairing small RNA Spot 42. Mol Microbiol 117:32–53.

65. Yu D, Ellis HM, Lee EC, Jenkins NA, Copeland NG, Court DL. 2000. An efficient recombination system for chromosome engineering in Escherichia coli. Proc Natl Acad Sci U S A 97:5978–83.

66. Court DL, Swaminathan S, Yu D, Wilson H, Baker T, Bubunenko M, Sawitzke J, Sharan SK. 2003. Mini-lambda: a tractable system for chromosome and BAC engineering. Gene 315:63–69.

67. Sharan SK, Thomason LC, Kuznetsov SG, Court DL. 2009. Recombineering: a homologous recombination-based method of genetic engineering. Nat Protoc 4:206–23.

68. Court DL, Swaminathan S, Yu D, Wilson H, Baker T, Bubunenko M, Sawitzke J, Sharan SK. 2003. Mini-lambda: a tractable system for chromosome and BAC engineering. Gene 315:63–9.

69. Thomason LC, Costantino N, Court DL. 2007. E. coli genome manipulation by P1 transduction. Curr Protoc Mol Biol Chapter 1:1 17 1-1 17 8.

70. Chung CT, Niemela SL, Miller RH. 1989. One-step preparation of competent *Escherichia coli*: transformation and storage of bacterial cells in the same solutions. PNAS USA 86:2172–2175.

71. Karlsson J, Eichner H, Loh E. 2023. Total Bacterial RNA Isolation and Northern Blotting Analysis. Methods Mol Biol 2674:73–85.

72. Ares M. 2012. Bacterial RNA isolation. Cold Spring Harb Protoc 2012:1024–7.

73. Masse E, Gottesman S. 2002. A small RNA regulates the expression of genes involved in iron metabolism in *Escherichia coli*. Proc Natl Acad Sci U S A 99:4620–5.

74. Rio DC. 2015. Northern blots: capillary transfer of RNA from agarose gels and filter hybridization using standard stringency conditions. Cold Spring Harb Protoc 2015:306–13.

75. Thibodeau SA, Fang R, Joung JK. 2004. High-throughput beta-galactosidase assay for bacterial cell-based reporter systems. Biotechniques 36:410–5.

76. London LY, Aubee JI, Nurse J, Thompson KM. 2021. Post-transcriptional regulation of rseA by small RNAs *ryhB* and *fnrS* in *Escherichia coli*. Front Mol Biosci 8:668613.

77. Jorgensen MG, Thomason MK, Havelund J, Valentin-Hansen P, Storz G. 2013. Dual function of the McaS small RNA in controlling biofilm formation. Genes Dev 27:1132–1145.

78. Luo H, Wei J, Wu S, Zheng Q, Zhang N, Chen P. 2023. Exploring CircRNA N6-methyladenosine in human rheumatoid arthritis: hyper-methylated hsa_circ_0007259 as a potential biomarker and its involvement in the hsa_circ_0007259/hsa_miR-21-5p/STAT3 axis. Int Immunopharmacol 124:110938.

79. He T, Hia H, Chen B, Duan Z, Huang C. 2023. m6A Writer mettl3-mediated lncRNA linc01125 prevents the malignancy of papillary thyroid cancer. Crit Rev Immunol 43:43–52.

80. Ma L, Ma Q, Deng Q, Zhou J, Zhou Y, Wei Q, Huang Z, Lao X, Du P. 2023. N7-methylguanosine-related miRNAs predict hepatocellular carcinoma prognosis and immune therapy. Aging (Albany NY) 3.

81. Chan CT, Deng W, Li F, DeMott MS, Babu IR, Begley TJ, Dedon PC. 2015. Highly Predictive Reprogramming of tRNA Modifications Is Linked to Selective Expression of Codon-Biased Genes. Chem Res Toxicol 28:978–88.

82. Deng W, Babu IR, Su D, Yin S, Begley TJ, Dedon PC. 2015. Trm9-Catalyzed tRNA Modifications Regulate Global Protein Expression by Codon-Biased Translation. PLoS Genet 11:e1005706.

83. de Crecy-Lagard V, Boccaletto P, Mangleburg CG, Sharma P, Lowe TM, Leidel SA, Bujnicki JM. 2019. Matching tRNA modifications in humans to their known and predicted enzymes. Nucleic Acids Res 47:2143–2159.

84. Helm M, Alfonzo JD. 2014. Posttranscriptional RNA Modifications: Playing Metabolic Games in a Cell’s Chemical Legoland. Chemistry and Biology 21:174–185.

85. Gu C, Begley TJ, Dedon PC. 2014. tRNA modifications regulate translation during cellular stress. FEBS Lett 588:4287–96.

86. Dedon PC, Begley TJ. 2014. A system of RNA modifications and biased codon use controls cellular stress response at the level of translation. Chem Res Toxicol 27:330–7.

87. Pollo-Oliveira L, Davis NK, Hossain I, Ho P, Yuan Y, Salguero Garcia P, Pereira C, Byrne SR, Leng J, Sze M, Blaby-Haas CE, Sekowska A, Montoya A, Begley T, Danchin A, Aalberts DP, Angerhofer A, Hunt J, Conesa A, Dedon PC, de Crecy-Lagard V. 2022. The absence of the queuosine tRNA modification leads to pleiotropic phenotypes revealing perturbations of metal and oxidative stress homeostasis in *Escherichia coli* K12. Metallomics 14.

88. Chionh YH, McBee M, Babu IR, Hia F, Lin W, Zhao W, Cao J, Dziergowska A, Malkiewicz A, Begley TJ, Alonso S, Dedon PC. 2016. tRNA-mediated codon-biased translation in mycobacterial hypoxic persistence. Nat Commun 11:13302.

89. Lampi M, Gregorova P, Qasim MS, AAhlblad NCV, Sarin LP. 2023. Bacteriophage infection of the marine bacterium *shewanella glacialimarina* induces dynamic change in tRNA modifications. Microorganisms 11:355.

90. Chionh YH, McBee M, Babu IR, Hia F, Lin W, Zhao W, Cao J, Dziergowska A, Malkiewicz A, Begley TJ, Alonso S, Dedon PC. 2016. tRNA-mediated codon-biased translation in mycobacterial hypoxic persistence. Nat Commun 7:13302.

91. Nakayashiki T, Inokuchi H. 1998. Novel temperature-sensitive mutants of *Escherichia coli* that are unable to grow in the absence of wild type tRNA_6_^Leu^. J Bacteriol 180:2931–2935.

92. Renda A, Poly S, Lai YJ, Pannuri A, Yakhnin H, Potts AH, Bevilacqua PC, Romeo T, Babitzke P. 2020. CsrA-Mediated Translational Activation of ymdA Expression in Escherichia coli. mBio 11.

93. Vakulskas CA, Leng Y, Abe H, Amaki T, Okayama A, Babitzke P, Suzuki K, Romeo T. 2016. Antagonistic control of the turnover pathway for the global regulatory sRNA CsrB by the CsrA and CsrD proteins. Nucleic Acids Res 44:7896–910.

94. Kitagawa R, Mitsuki H, Okazaki T, Ogawa T. 1996. A novel DnaA protein-binding site at 94.7 min on the Escherichia coli chromosome. Mol Microbiol 19:1137–47.

95. Maciag A, Peano C, Pietrelli A, Egli T, De Bellis G, Landini P. 2011. In vitro transcription profiling of the sigmaS subunit of bacterial RNA polymerase: re-definition of the sigmaS regulon and identification of sigmaS-specific promoter sequence elements. Nucleic Acids Res 39:5338–55.

96. Tsui HC, Leung HC, Winkler ME. 1994. Characterization of broadly pleiotropic phenotypes caused by an hfq insertion mutation in *Escherichia coli* K-12. Mol Microbiol 13:35–49.

97. Brown L, Elliot T. 1996. Efficient translation of the RpoS sigma factor in *Salmonella typhimurium* requires host factor I, an RNA-binding protein encoded by the *hfq* gene. J Bacteriol 178:3763–3770.

98. Brown L, Elliot T. 1997. Mutations that increase expression of the *rpoS* gene and decrease its dependence on *hfq* function in *Salmonella typhimurium*. J Bacteriol 179:656–662.

99. Zhang A, Altuvia S, Tiwari A, Argaman L, Hengge-Aronis R, Storz G. 1998. The OxyS regulatory RNA represses *rpoS* translation and binds the Hfq (HF-I) protein. EMBO J 17:6061–6068.

100. Sledjeski DD, Whitman C, Zhang A. 2001 Hfq is necessary for regulation by the untranslated RNA DsrA. J Bacteriol 183:1997–2005.

101. Majdalani N, Chen S, Murrow J, St John K, Gottesman S. 2001. Regulation of RpoS by a novel small RNA: the characterization of RprA. Mol Microbiol 39:1382–94.

102. Baker CS, Eory LA, Yakhnin H, Mercante J, Romeo T, Babitzke P. 2007. CsrA inhibits translation initiation of *Escherichia coli hfq* by binding to a single site overlapping the shine-dalgarno aequence. J Bacteriol 189:5472–5481.

